# Phosphorylation of LXRα impacts atherosclerosis regression by modulating monocyte/macrophage trafficking

**DOI:** 10.1101/363366

**Authors:** Elina Shrestha, Maud Voisin, Tessa J. Barrett, Hitoo Nishi, David J. Cantor, Maryem A. Hussein, Gregory David, Inés Pineda-Torra, Edward A. Fisher, Michael J. Garabedian

## Abstract

LXRα activation in macrophages enhances regression of atherosclerotic plaques in mice by regulating genes crucial for cholesterol efflux, cell motility and inflammation. Diabetes, however, impairs plaque regression in mice. LXRα is phosphorylated at serine 198 (pS198), which affects the expression of genes controlling inflammation, lipid metabolism and cell movement. We hypothesize that LXRα function is affected by hyperglycemia through changes in LXRα pS198. Indeed, macrophages cultured in diabetes relevant high glucose versus normal glucose display alterations in LXR-dependent gene expression and increased LXRα pS198. We therefore examined the consequence of disrupting LXRα phosphorylation (S196A in mouse LXRα) during regression of atherosclerosis in normal and diabetic mice. We find that phosphorylation deficient LXRα S196A reduces macrophage retention in plaques in diabetes, which is predicted to be anti-atherogenic and enhance plaque regression. However, this favorable effect on regression is masked by increased monocyte infiltration in the plaque attributed to leukocytosis in LXRα S196A mice. RNA-seq of plaque macrophages from diabetic S196A mice shows increased expression of chemotaxis and decreased expression of cell adhesion genes, consistent with reduced macrophage retention by LXRα S196A. Thus, the non-phosphorylated form of LXRα precludes macrophage retention in the plaque. Our study provides the first evidence for a physiological role of LXRα phosphorylation in modulating atherosclerosis regression. Compounds that prevent LXRα phosphorylation or ligands that induce the conformation of non-phosphorylated LXRα may selectively enhance macrophage emigration from atherosclerotic plaques.

## INTRODUCTION

Atherosclerosis is an inflammatory disease characterized by lipid-rich plaques in the subendothelial space of arterial walls that contain cholesterol-loaded macrophage foam cells (1). If left untreated, a plaque can rupture leading to myocardial infarction or stroke. A major risk factor for atherosclerosis is diabetes (2), the incidence of which continues to rise (3).

Normalization of cholesterol levels has been shown to promote macrophage egress and the regression of atherosclerotic plaques in some mouse models [reviewed in (4)]. Mice deficient in Liver X Receptors (LXRs) displayed impaired regression through a defect in the emigration of plaque macrophages after hypercholesterolemia was reversed (5). LXRs, LXRα and LXRβ, belong to the nuclear receptor superfamily of DNA-binding transcription factors and are activated by binding oxysterol cholesterol metabolites and cholesterol precursors, and synthetic agonists (6-10). Increased intracellular cholesterol levels in macrophages activate LXRs, which induces expression of several genes important in cholesterol efflux and reverse cholesterol transport. Previously our lab showed that human LXRα is phosphorylated at serine 198 (which corresponds to serine 196 in mouse LXRα) by Casein kinase 2 (CK2) in cultured macrophages (11). This modification has been shown to selectively regulate LXR target gene expression, suggesting that it might impact LXR signaling in macrophages in the atherosclerotic plaque (12). In mice, increased LXRα phosphorylation at S196 (hereafter LXRα pS196 in mouse and LXRα pS198 in human) in plaque macrophages is associated with atherosclerosis progression whereas reduced LXRα pS196 is associated with regression (12). However, the causal contribution of LXRα phosphorylation in atherosclerosis *in vivo* has not been examined.

Plaque regression upon reductions in LDL cholesterol levels is significantly impaired in diabetic vs. non-diabetic mice (13). This is consistent with clinical data showing that diabetes significantly hampers the regression of human atherosclerosis, despite the equivalent drop in lipid levels (14). Regression in diabetic mice is associated with defective reduction of plaque CD68+ cells, suggesting increased macrophage retention or increased entry of monocytes despite lipid lowering (13, 15). In a recent study we found that chronic high glucose significantly compromised LXR-dependent gene activation relative to normoglycemia in cultured bone marrow derived macrophages (16). Because of increased phosphorylation of LXRα at S196 in progressing plaques (12), we hypothesized that LXR function is impaired in diabetic mice through the same modification and contributes to the impaired regression.

In this report, we test this hypothesis by interrogating the impact of high glucose concentrations on LXRα pS196 in macrophages. In addition, to better understand the biological function of LXRα phosphorylation, we generated a mouse in which the serine 196 codon has been changed to alanine (S196A), to test the physiological relevance of LXRα pS196 in the regression of atherosclerosis in normoglycemic and diabetic settings. Remarkably, the genetic prevention of LXRα phosphorylation reduces macrophage retention in plaques during regression of atherosclerosis in diabetes. Underlying this reduction in macrophage retention are changes in gene expression by the non-phosphorylated LXRα that promote chemotaxis and reduce cell adhesion that contribute to this important anti-atherogenic phenotype.

## RESULTS

### Hyperglycemia influences LXR-dependent gene expression and enhances LXRα phosphorylation in macrophages

Atherosclerosis regression upon lipid lowering is hampered in hyperglycemic mice (13, 17). Given the importance of LXRs in atherosclerosis regression (5), we hypothesized that hyperglycemia influences receptor function. Thus, we performed gene expression-profiling on RAW 264.7 murine macrophages stably expressing human LXRα cultured in diabetes-relevant high glucose (25 mM D-glucose) or normal glucose (5.5 mM D-glucose, supplemented with inactive L-glucose to control for osmolality) with and without LXR agonists (Supplemental Figure 1A). This allowed us compare the impact of glucose on the transcriptional response of macrophages to LXR activation. High compared to normal glucose significantly modulated 1,297 genes; 647 genes exhibited enhanced and 650 displayed reduced expression (>1.5 fold; p-value <0.05). Interestingly, of the 128 genes that were induced upon ligand treatment, 48% of genes (61 out of 128) were modulated by glucose: 31 genes were further induced, whereas 30 genes had reduced ligand-dependent induction in high compared to normal glucose. On the other hand, of the 48 genes that were repressed by high glucose, 42% of genes (20 out of 48) were sensitive to the increase in glucose concentration: 11 genes were further repressed whereas 9 genes had less repression with ligand in high compared to normal glucose (Figure 1A). This supported our hypothesis that glucose has a significant effect on LXR-dependent gene expression. Importantly, the gene expression changes affected by high glucose were linked to molecular and cellular functions in macrophages including cell death and survival, cell movement, cell growth and lipid metabolism (Figure 1B). High glucose also appeared to control inflammatory signaling, atherosclerosis signaling as well as LXR activation pathways (Supplemental Figure 1B). Thus, glucose-regulated changes in LXR-dependent gene expression in macrophages affects key determinants of atherosclerosis.

**Figure 1.**
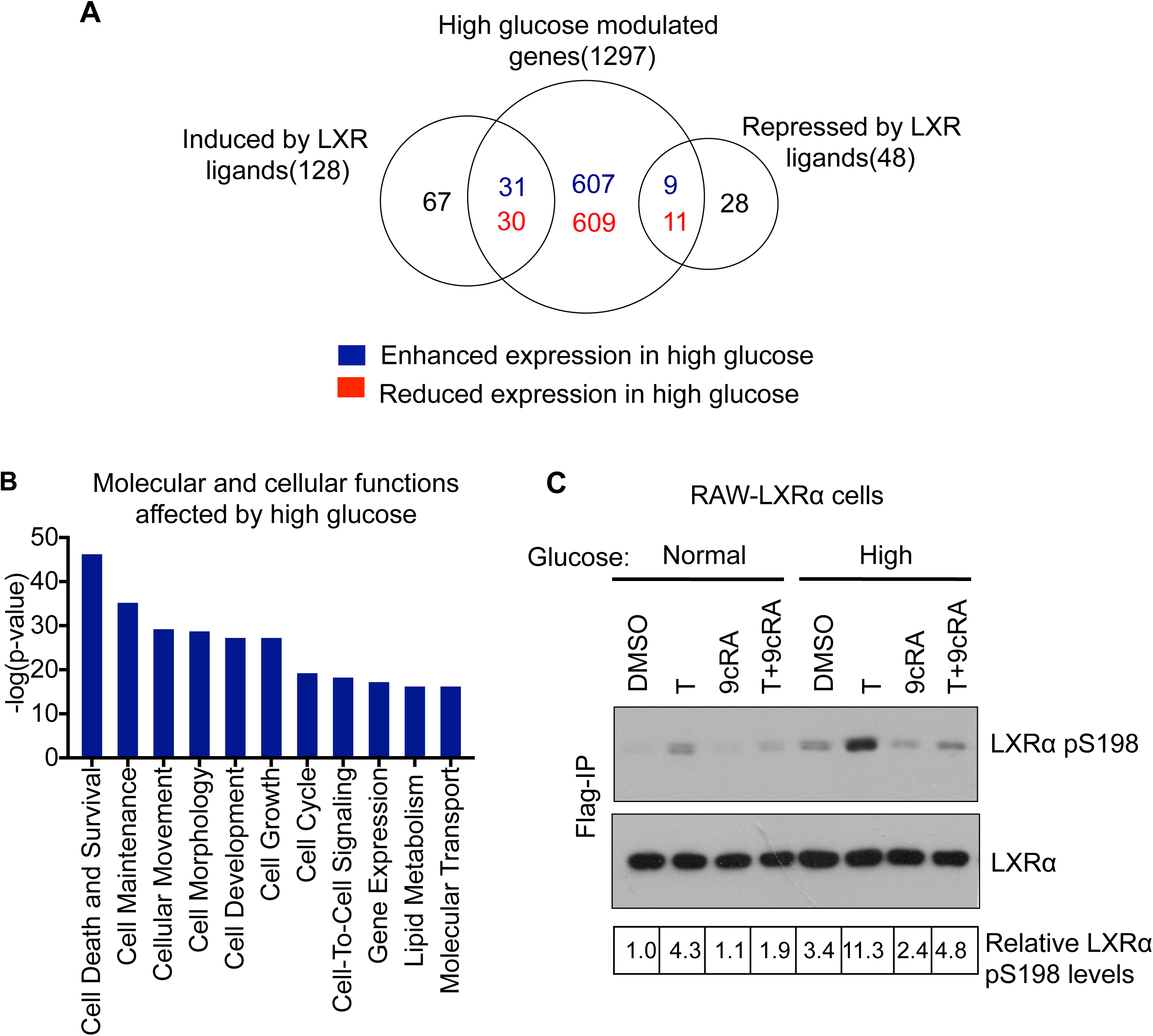
Hyperglycemia influences LXR signaling and enhances LXRα S198 phosphorylation. A) RAW 264.7 macrophages stably expressing human LXRα (RAW-LXRα) were cultured under high glucose (25mM D-glucose) or normal glucose (5.5mM D-glucose supplemented with L-glucose as an osmotic control). Macrophages were treated with LXR/RXR ligands (5uM T0901317 +1uM 9cisRA) for 4 hours. Total RNA was isolated and gene expression was assessed using the Mouse Genome 430 2.0 AffyMetrix Expression Array. Four independent biological replicates were performed per condition. Venn diagram showing overlap between genes affected (induced and repressed > 1.5 fold; p < 0.05) by high compared to normal glucose in the absence and presence of ligand treatment. The numbers of genes downregulated are shown in red and upregulated are shown in blue in high compared to normal glucose. B) Molecular and cellular functions of genes regulated by high versus normal glucose under ligand treated conditions predicted by IPA. C) RAW-LXRα cells were cultured in normal and high glucose, and treated with vehicle control (DMSO), the LXR ligand (5 μM T0901317; T), RXR ligand (1 μM 9-cisRA; 9cRA) or the combination of both for 4 h. LXRα was immunoprecipitated with anti-Flag antibody and LXRα protein and S198 phosphorylation were detected using an LXRα phospho-S198 and total LXRα antibodies. The Western blot was quantitated using ImageJ software, and the ratio of LXRα pS198 to LXRα is shown for each sample with DMSO treated sample in normal glucose set to 1.

We next sought to interrogate how hyperglycemia influences LXR signaling. LXRα phosphorylation at S198 is known to modulate its transcriptional activity at a subset of target genes, and promotes pro-inflammatory gene expression while suppressing anti-inflammatory gene expression in macrophages (11). We therefore explored the possbility that high glucose affected LXRα function through changes in LXRα phosphorylaiton. RAW 264.7 cells stably expressing human LXRα cultured either under normal or diabetes relevant high glucose were examined for effects on LXRα S198 phosphorylation. Cells were treated with DMSO as the vehicle control, T0901317, 9-cisRA or T0901317 and 9-cisRA. This combination of ligands reduces LXRα pS198 induced by T0901317 (11). Consistent with our hypothesis, LXRα pS198 is greatly increased under high compared to normal glucose under basal and ligand-treated conditions, suggesting a glucose-dependent regulation of LXR phosphorylation (Figure 1C).

### Expression of phosphorylation deficient LXRα S196A in mice does not change plasma lipid and glucose levels

Next we determined whether LXRα phosphorylation affected macrophage function during atherosclerosis regression in diabetes. To do this, we used an inducible LXRα S196A phosphorylation deficient knock-in mouse (Supplemental Figure 2). We employed this strategy so plaques could develop normally under conditions of LXRα WT and then be switched to LXRα S196A to isolate the effect of LXR phosphorylation on regression (see Figure 2A for study protocol). Mice homozygous for LXRα S196A floxed allele (S196A^FL/FL^) (18) with or without a tamoxifen inducible CRE recombinase (RosaCreER^T2^;CRE+ or CRE-) were used as donors for bone marrow (BM) transplantation into recipient Reversa mice (19). This is an *Ldlr*^−/−^ model where the hyperlipidemia can be reversed after conditional inactivation of microsomal triglyceride transfer protein (*Mttp*) by pIpC treatment (Figure 2A). Reversa mice were fed a western diet for 16 weeks to allow plaques to develop.

**Figure 2.**
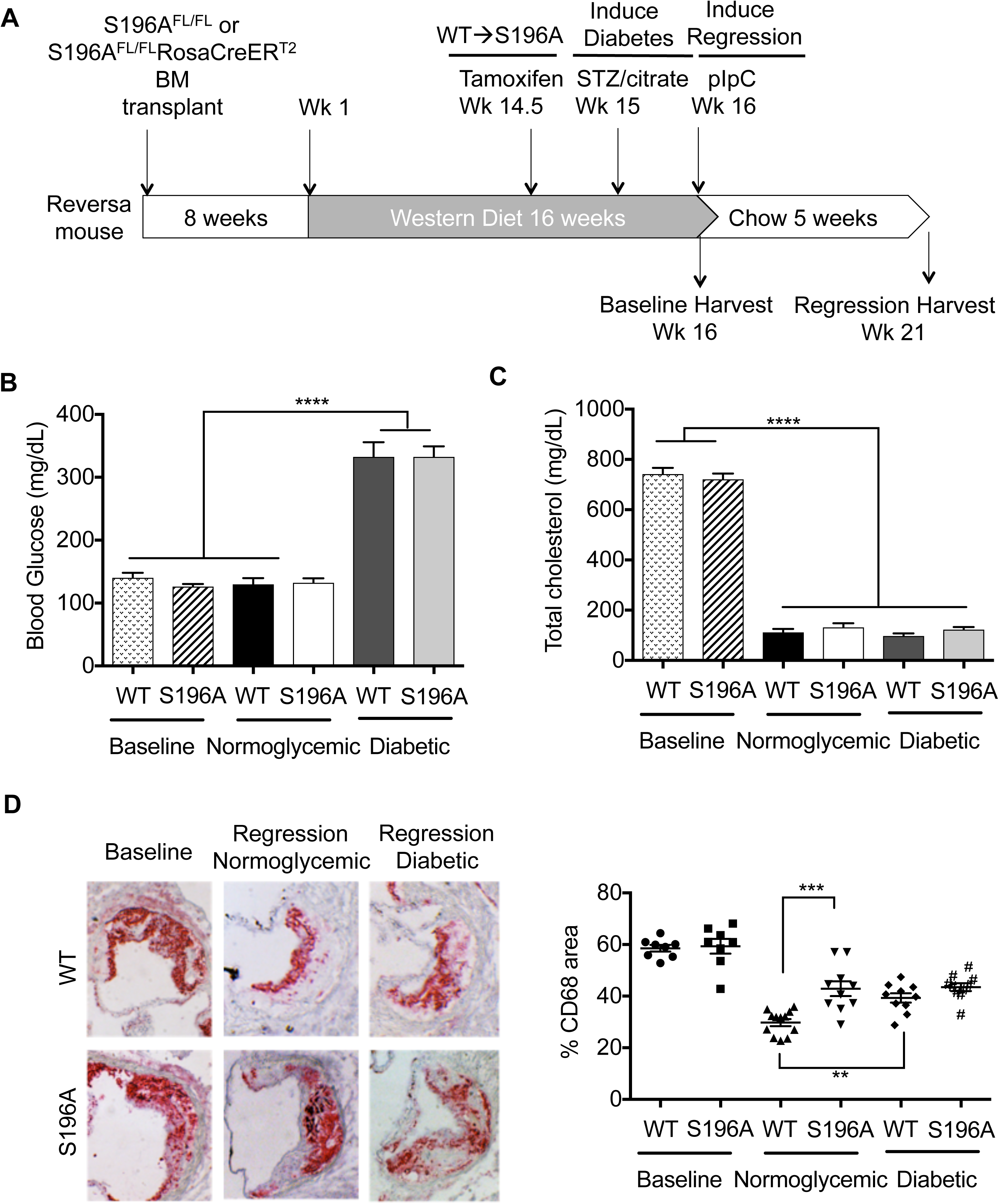
LXRα S196A expression increases plaque macrophage content in normoglycemic mice but not in diabetic mice. A) Experimental design to test the effects of LXRα pS196 in atherosclerosis regression under normoglycemia and diabetes. B) Blood glucose levels were measure before harvesting the mice. C) Blood was collected from the mice at the time of harvest and total cholesterol was measured by using colorimetric assays. D) Aortic roots from baseline and regression groups were sectioned, fixed, and stained for CD68 to measure total macrophage content. Representative images of CD68 immunostaining are shown for each group (left panel). The percent area occupied by CD68+ cells of the plaque area was quantified using Image Pro Plus Software. Data (mean ± SEM) (n≥8) were analyzed using one-way ANOVA followed by Tukey’s multiple comparison test. P < 0.05 values were considered to be significant, ^∗^P < 0.05, ^∗∗^P < 0.005, ^∗∗∗^P < 0.0005 and ^∗∗∗∗^P < 0.00005. WT; S 196A^FL/FL^ CRE-recipient Reversa mice; SA: S196A^FL/FL^ CRE+ recipient Reversa mice.

Mice were separated into six groups: two groups of baseline mice (LXRα S196A^FL/FL^ CRE- and LXRα S196A^FL/FL^ CRE+), and four regression groups; 1) LXRα WT (S196A^FL/FL^ CRE-) and normoglycemia, 2) LXRα S196A (S196A^FL/FL^ CRE+) and normoglycemia; 3) LXRα WT (S196A^FL/FL^ CRE-) and diabetic; 4) LXRα S196A (S196A^FL/FL^ CRE+) and diabetic. Tamoxifen was administered at week 14.5 to all groups and this induced recombination of loxP sites in the CRE expressing mice resulting in the replacement of the LXRα WT with the LXRα S196A allele. The regression/diabetic mice were administered an intraperitoneal injection of Streptozotocin (STZ) for 5 days preceding pIpC treatment so that they became hyperglycemic.

The reconstitution of bone marrow after transplantation in Reversa mice was efficient: 90% of the circulating monocytes were derived from the newly transplanted bone marrow (Supplemental Figure 3). The CRE-mediated recombination at the S196A allele upon tamoxifen treatment was also effective. Notably, the atherosclerotic plaques in the aortic roots of mice expressing LXRα S196A (S196A^FL/FL^ CRE+ BM recipients) versus those that express LXRα WT (S196A^FL/FL^ CRE-BM recipients) showed markedly reduced LXRα pS196 staining in plaques, without changes in total LXRα abundance (Supplemental Figure 4 and 5). This is indicative of an efficient switch to LXRα S196A protein in the plaque.

The LXRα S196A (S196A^FL/FL^ CRE+) BM recipients had similar glucose levels and lipid profiles (cholesterol, triglycerides and HDL cholesterol) compared to their LXRα WT (S196A^FL/FL^ CRE-) BM recipient counterparts, suggesting that LXRα S196A expression during the 5 weeks of regression did not affect these parameters (Figure 2B and 2C and Supplemental Figure 6A and 6B). As expected, the glucose levels were dramatically higher in STZ-treated mice than in control mice (Figure 2B). Similar reductions in cholesterol and triglyceride levels were observed in normal and diabetic mice upon reversal of hyperlipidemia (Figure 2C and Supplemental Figure 6A).

### Expression of LXRα S196A increases plaque macrophage content in normoglycemic but not diabetic mice

To determine the effect of LXRα S196A on atherosclerosis regression, we measured plaque area and their total macrophage content by quantifying the percent of the macrophage marker CD68 stained area. Both groups of baseline mice had similar macrophage contents in their plaques since the plaques developed largely under LXRα WT conditions (Figure 2D). After 5 weeks of reduction in plasma cholesterol levels, we observed reduced macrophage content in the normoglycemic mice expressing LXRα WT indicative of atherosclerosis regression. However, in mice expressing LXRα S196A regression was impaired as evidenced by significantly less reduction in CD68 positive cell content compared to LXRα WT groups under normoglycemic conditions (Figure 2D). Consistent with previous findings, diabetic mice with LXRα WT also showed impaired plaque regression (13, 15). Interestingly, unlike the normoglycemic setting, LXRα S196A expression did not alter plaque macrophage content under diabetic conditions: regression was similarly impaired in diabetic mice expressing LXRα WT and LXRα S196A. As shown before, the total plaque area in aortic roots of the baseline and regression groups as well as normoglycemic and diabetic groups remained unchanged (Supplemental Figure 7) (13, 15). Collagen content in the plaques among all the regression groups was also unchanged (not shown). Thus, LXRα S196A expression increased plaque macrophage burden and impaired regression in normoglycemic mice but did not affect plaque regression in diabetic mice.

### LXRα pS196 and diabetes affect apoptosis, necrotic area, and efferocytosis

In advanced lesions, macrophages undergo apoptosis, and this can affect macrophage content within a plaque (24). Therefore, we measured apoptosis in the plaques by staining for cleaved caspase 3. We found increased apoptosis in LXRα S196A plaques under normoglycemic conditions compared to LXRα WT (Figure 3A). Relative to WT normoglycemia, diabetes also increased apoptosis in the plaque of LXRα WT expressing mice, which was not further increased by LXRα S196A.

**Figure 3.**
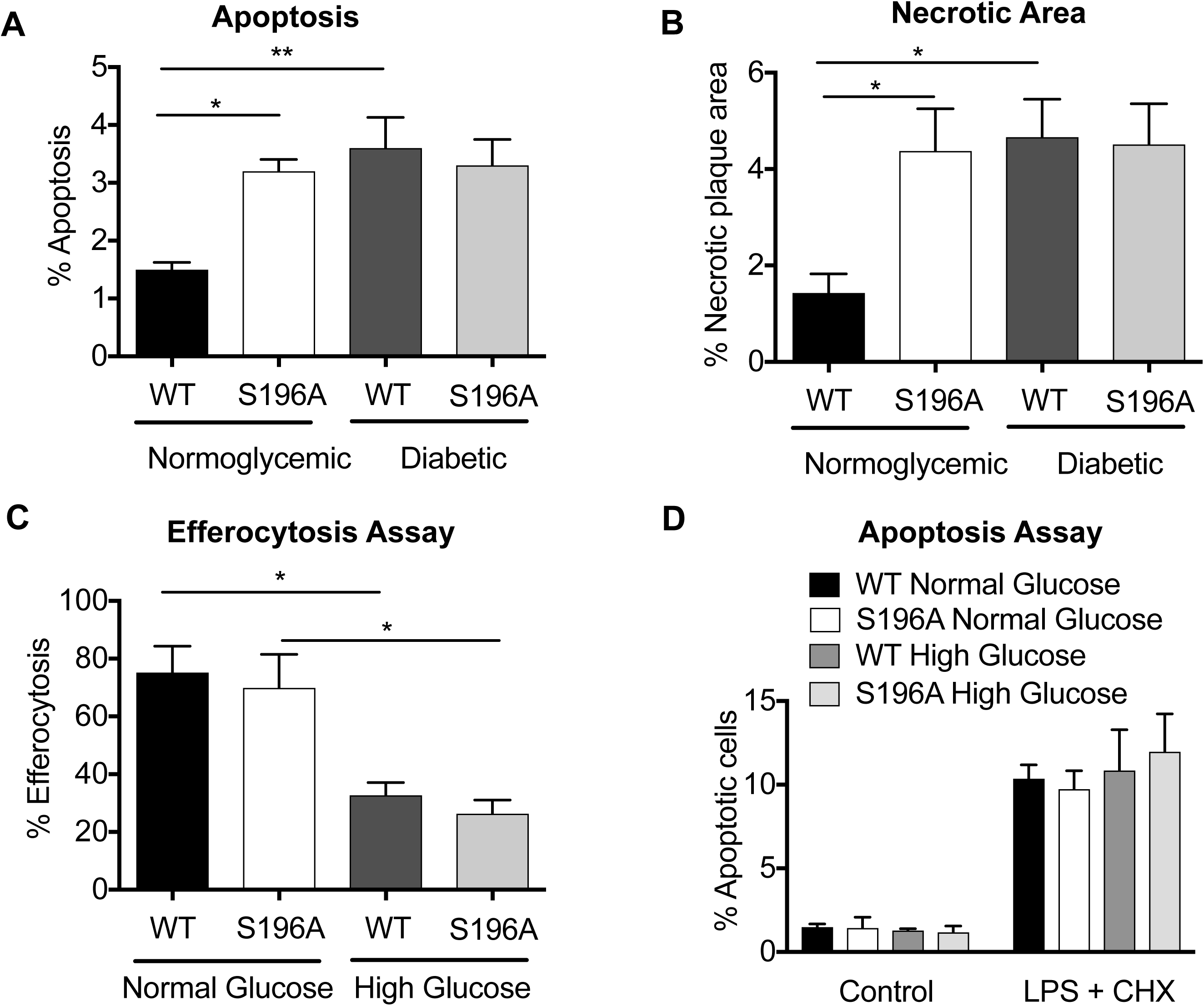
LXRα S196A expression increases plaque necrotic area and apoptosis in normoglycemic mice but not in diabetic mice. The percent necrotic area of plaques from the regression groups are shown. Data (mean ± SEM; n≥6) were analyzed using one-way ANOVA followed by Tukey’s multiple comparison test. P < 0.05 values were considered to be significant, ^∗^P < 0.05, ^∗∗^P < 0.005 and ^∗∗∗^P < 0.005. B) Aortic roots from the regression groups were stained for cleaved caspase 3 to detect cells undergoing apoptosis. The number of apoptotic cells per section were counted and precent apoptotic cells in the plaque are shown. Data (mean ± SEM; n≥7) were analyzed as above. C) Apoptotic Jurkat cells were added to peritoneal macrophages that express LXRα WT or LXRα S196A cultured under normal and high glucose conditions, and efferocytosis was quantified and expressed as the percent of cells having ingested apoptotic cells. D) Peritoneal macrophages expressing LXRα WT or LXRα S196A under normal glucose and high glucose were treated with LPS and cyclohexamide to induce apoptosis for 4 hours and % apoptosis was measured. The experiments in C and D were performed three times with similar results. Each experiment was performed in triplicate, and representative experiments are shown. The error bars represent SEM. Significance was determined as above;. ^∗^P < 0.05.

When the apoptotic cells are not efficiently cleared by macrophages in a process called efferocytosis, they promote the formation and expansion of a necrotic core that creates plaques vulnerable to rupture and thrombosis (3). We measured necrotic area by quantifying the acellular areas within the plaques (15), and found that plaques from LXRα S196A-expressing mice contained significantly larger necrotic areas compared to LXRα WT mice under normoglycemia (Figure 3B). Diabetes also significantly increased necrotic areas in wild type mice consistent with previous reports (2), whereas LXRα S196A did not further exacerbate plaque necrosis in diabetic mice.

Since regressing plaques in some groups of mice were associated with significantly higher necrotic areas compared to others, this suggested inadequate or defective clearance of dying or dead cells by efferocytosis. To test whether hyperglycemia and/or phosphorylation of LXRα at S196 affected efferocytosis, we assayed the ability of peritoneal macrophages expressing LXRα WT and LXRα S196A cultured in normal and high glucose to efferocytose labeled apoptotic Jurkat cells. The uptake of apoptotic cells (% efferocytosis) was significantly reduced by culturing cells in high glucose, and independent of LXRα S196 phosphorylation (Figure 3C). This would help to explain the larger necrotic core in the diabetic atherosclerotic plaques.

Since LXRα S196 phosphorylation in macrophages had no effect on efferocytosis, this suggested that higher plaque necrotic area in mice expressing LXRα S196A compared to LXRα WT under normoglycemic condition was due to other factors, such as increased apoptosis. We therefore examined the propensity of peritoneal macrophages expressing LXRα WT and LXRα S196A to undergo apoptosis induced by LPS and cyclohexamide under normal glucose and high glucose conditions. Similar levels of apoptosis were observed in LXRα WT and LXRα S196A-expressing cells under hyperglycemic and normoglycemic conditions (Figure 3D). Therefore, differences in the content of apoptotic cells in the plaque were likely not due to cell intrinsic changes in apoptotic activity of macrophages between LXRα WT and LXRα S196A, but rather, the increased macrophage accumulation in the plaque.

### Loss of LXRα S196 phosphorylation increased monocyte recruitment and reversed diabetes-associated increased macrophage retention

We next investigated whether there were alterations in monocyte/macrophage trafficking in the plaque that could be responsible for differences in macrophage content among the regression groups, using methods we have previously used for this purpose (15, 20). To measure monocyte recruitment, mice were injected with fluorescent latex beads 24 hours before harvesting the aortic roots, and the number of beads in the plaque was quantified (Figure 4A). Monocyte labeling was not affected by either diabetes or LXRα S196A expression suggesting that the phagocytic ability of circulating monocytes was similar in all groups (Supplementary Figure 8A). Diabetes increased monocyte recruitment in mice expressing LXRα WT (Figure 4B), consistent with previous studies (15). Interestingly, mice expressing LXRα S196A also displayed significantly increased monocyte recruitment under both normoglycemic and diabetic conditions. This suggests that LXRα S196A expressing mice have greater CD68+ plaque area under normoglycemia due to enhanced monocyte recruitment. Although LXRα S196A expression increased recruitment of monocytes under diabetes, however, it did not alter the CD68+ plaque area compared to LXRα WT. This indicates that LXRα S196A expression is affecting another pathway that compensates for increased monocyte recruitment. For example, LXRα S196A expression may inhibit macrophage retention/emigration under diabetes.

**Figure 4.**
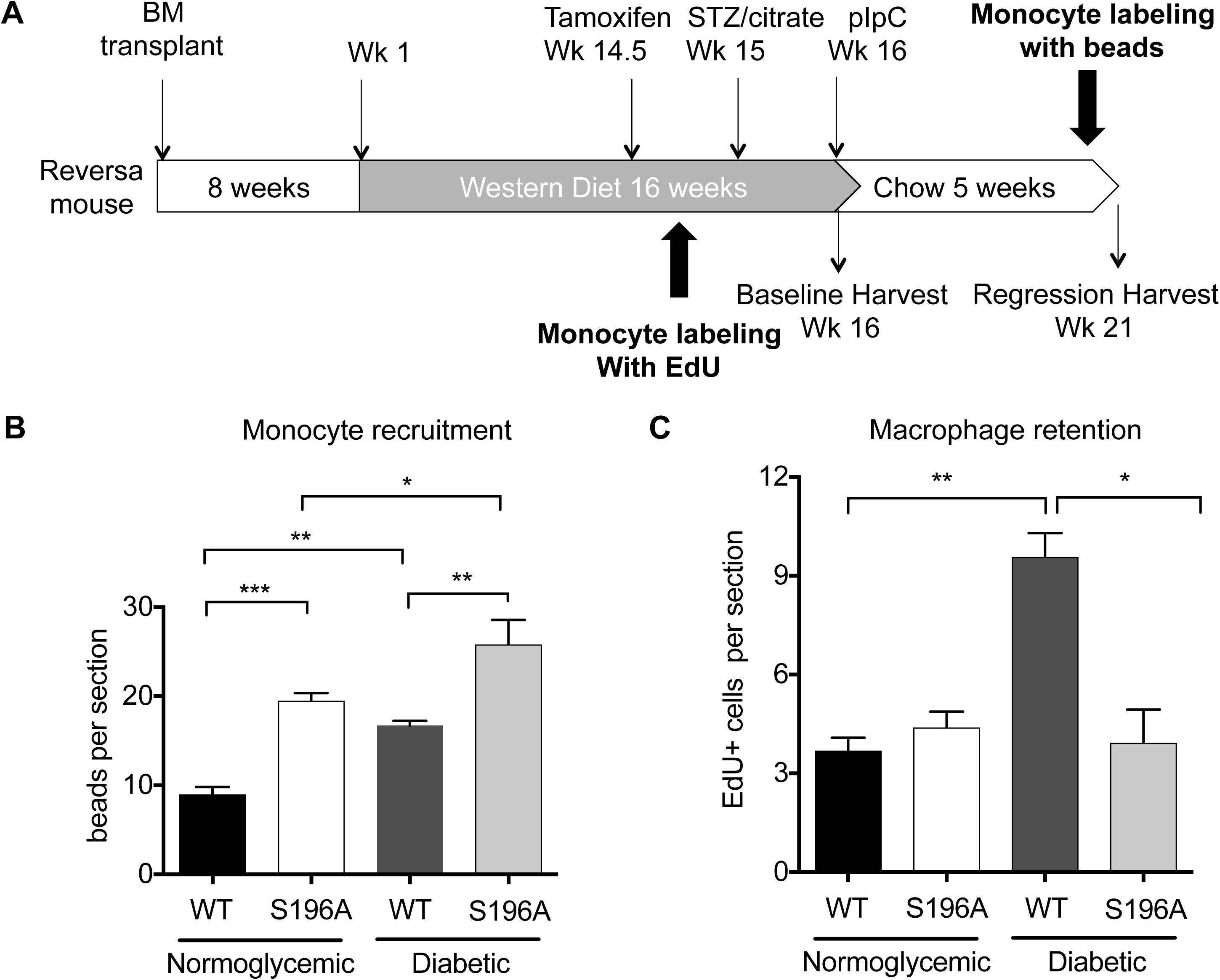
LXRα S196A expression and diabetes influence monocyte/macrophage trafficking in the plaque. A) Reversa mice were injected with EdU 6 days before the end of western diet feeding to measure macrophage retention in the plaque and with fluorescent latex beads 24 h before sacrificing the mice to assess monocyte recruitment. B) To measure monocyte recruitment, beads were counted in the aortic root using fluorescent microscopy. C) To assess macrophage retention the number of EdU positive cells were counted in aortic root from the regression groups. Data are expressed as mean ± SEM (n≥6 in each group). Data were analyzed using one-way ANOVA followed by Tukey’s multiple comparison test. P < 0.05 values were considered to be significant, ^∗^P < 0.05, ^∗∗^P < 0.005 and ^∗∗∗^P < 0.005.

To test whether LXRα S196A expression in macrophages under diabetes affects macrophage retention, mice were injected with EdU (5-ethynyl-2’-deoxyuridine) 6 days before the end of western diet feeding. EdU gets incorporated into DNA of proliferating progenitor cells, which give rise to circulating Ly6C^hi^ monocytes (Supplemental Figure 6B). After 24 hours of EdU injection ~50% of circulating Ly6C^hi^ monocytes were found to be labeled with EdU (Supplemental Figure 8B). We tracked the number of labeled cells that infiltrated into the plaques of WT and S196A baseline groups and did not find differences between the two groups (Supplemental Figure 8C). At the time of harvesting baseline groups, after 5 days of injection, EdU positive cells were not detected in circulation suggesting no new EdU positive cells would enter the plaques after this time point (Supplemental Figure 8B). Therefore, the number of EdU positive cells in the plaques after 5 weeks of lipid reduction would reflect macrophages retained in the plaque. We found fewer EdU positive macrophages in regression groups compared to the baseline consistent with cells emigrating out of the plaque over time (Figure 4C). We also found that diabetes is associated with increased macrophage retention in mice expressing LXRα WT: compared to normoglycemic LXRα WT mice, diabetic LXRα WT mice had approximately 50% more macrophages retained in their plaques (Figure 4C). Notably, LXRα S196A expression prevented the diabetes-induced increase in macrophage retention. This was independent of macrophage proliferation, since similar numbers of cells stained positive for Ki67, a marker of cell proliferation, among the regression groups (Supplemental Figure 9). Thus, LXRα S196A prevents macrophage retention in diabetes. Given that regression of atherosclerosis in Reversa mice has been shown to require egress of macrophages from the plaque (21), these findings suggest that LXRα S196A facilitates plaque regression by preventing macrophage accumulation in diabetes.

### LXRα S196A expressing mice have more circulating monocytes under regression conditions

The increased recruitment of monocytes in the LXRα S196A mice after lipid lowering could result from higher numbers of circulating leukocytes (22). In fact, we observed significantly higher levels of circulating monocytes and WBCs in LXRα S196A versus LXRα WT expressing mice in regression groups under both normoglycemic and diabetic conditions (Figure 5A). This likely explains the increase in the CD68+ cells in normoglycemic LXRα S196A versus LXRα WT-expressing mice during regression. This also suggests that LXRα pS196 plays a role in modulating circulating levels of WBCs.

**Figure 5.**
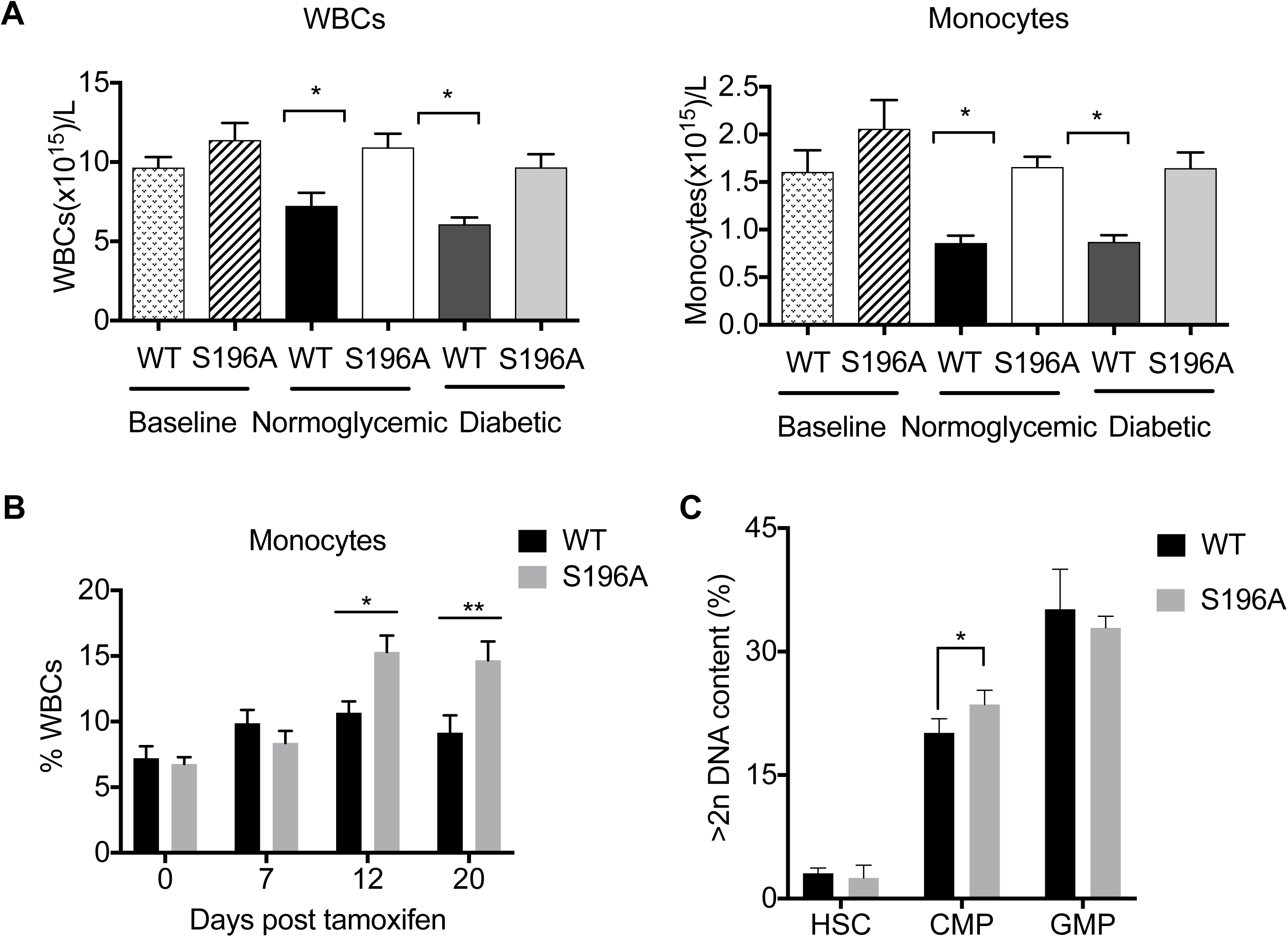
LXRα S196A expressing mice have more circulating monocytes and increased CMP proliferation during regression. A) Quantification of circulating monocytes and WBCs in mice. Data are expressed as mean ± SEM (n≥8 per group). Data were analyzed using one-way ANOVA followed by Tukey’s multiple comparison test. P < 0.05 values were considered to be significant, ^∗^P < 0.05, ^∗∗^P < 0.005 and ^∗∗∗^P < 0.005. B) S196A^FL/FL^Cre- (WT) and S196A^FL/FL^Cre+ (S196A) mice were treated with tamoxifen on Day 0. Blood monocytes were measured on Day 0, 7, 12 and 20 as percentage of WBCs was determined using Genesis Hematology system. C) Hematopoetic stem and progenitor cell populations in the bone marrow in G_2_M phase (cells with >2n DNA content) was measured by flow cytometry suing DAPI. CMP; common myeloid progenitor; GMP, granulocyte-macrophage progenitor; and HSC, hematopoetic stem cell.

### LXRα S196A increases CMP proliferation in bone marrow

Given that LXRα S196A expressing mice have higher monocytes compared to LXRα WT under both normoglycemic and diabetic regression conditions, we examined whether LXRα S196A affected circulating monocyte levels in the absence of dietary and blood glucose modulation. The monocyte counts in LXRα WT (S196A^FL/FL^ CRE-) and LXRα S196A (S196A^FL/FL^ CRE+) mice fed normal chow diet were measured before and after tamoxifen administration. Blood monocytes were monitored over 20 days. Induction of LXRα S196A expression led to increased blood monocytes 12 days after the first tamoxifen injection (Figure 5B). In contrast, monocytes were not affected in mice expressing LXRα WT after tamoxifen injection. At both 12 and 20-days post-tamoxifen treatment, mice that expressed LXRα S196A showed significantly higher levels of blood monocytes compared to those that expressed LXRα WT. This suggests that persistent LXRα S196A expression in the bone marrow of mice leads to an increase in blood monocytes.

To examine if hematopoietic progenitor cells were affected by LXRα S196A (23), we harvested bone marrow cells from normoglycemic and normolipidemic LXRα WT and LXRα S196A mice 20 days after recombination to the S196A allele. We analyzed the proliferation of monocyte progenitors, including hematopoietic stem cells (HSCs), common myeloid progenitors (CMPs) and granulocyte-monocyte progenitors (GMPs) by flow cytometry, as we have done before (15). We found that LXRα S196A expression increased the proliferation of the CMPs without affecting GMP or HSC populations (Figure 5C).

### RNA-seq analysis reveals diverse effects of LXRα S196 phosphorylation on gene expression of plaque macrophages under normoglycemic and diabetic settings

To gain insight into the mechanisms of changes in macrophage retention, we examined genome-wide expression changes from LXRα S196A and LXRα WT plaque macrophages from normoglycemic and diabetic mice by performing RNA sequencing from laser-captured CD68+ cells from plaques of regression groups. Under normoglycemic conditions, 447 genes (315 induced and 132 repressed) were modulated by LXRα S196A (>1.2 fold change, p<0.01) compared to expression under LXRα WT (Figure 6A and Supplementary Figure 10A). Under diabetic conditions, 212 genes (135 induced and 77 repressed) were modulated by LXRα S196A (Figure 6A and 6C). Intriguingly, the effects of LXRα S196 phosphorylation were highly dependent on the glucose environment because LXRα S196A regulated distinct sets of genes under normal and diabetic conditions (Figure 6B). Moreover, a significant number of genes that were upregulated by LXRα S196A under diabetes were downregulated by LXRα S196A under normoglycemia and those that were downregulated by LXRα S196A under diabetes were upregulated by LXRα S196A in normoglycemia (Table S1). This suggested that changes in glucose levels can switch the propensity of LXRα S196A from being a transcriptional activator to a repressor or vice-versa at certain genes.

**Figure 6.**
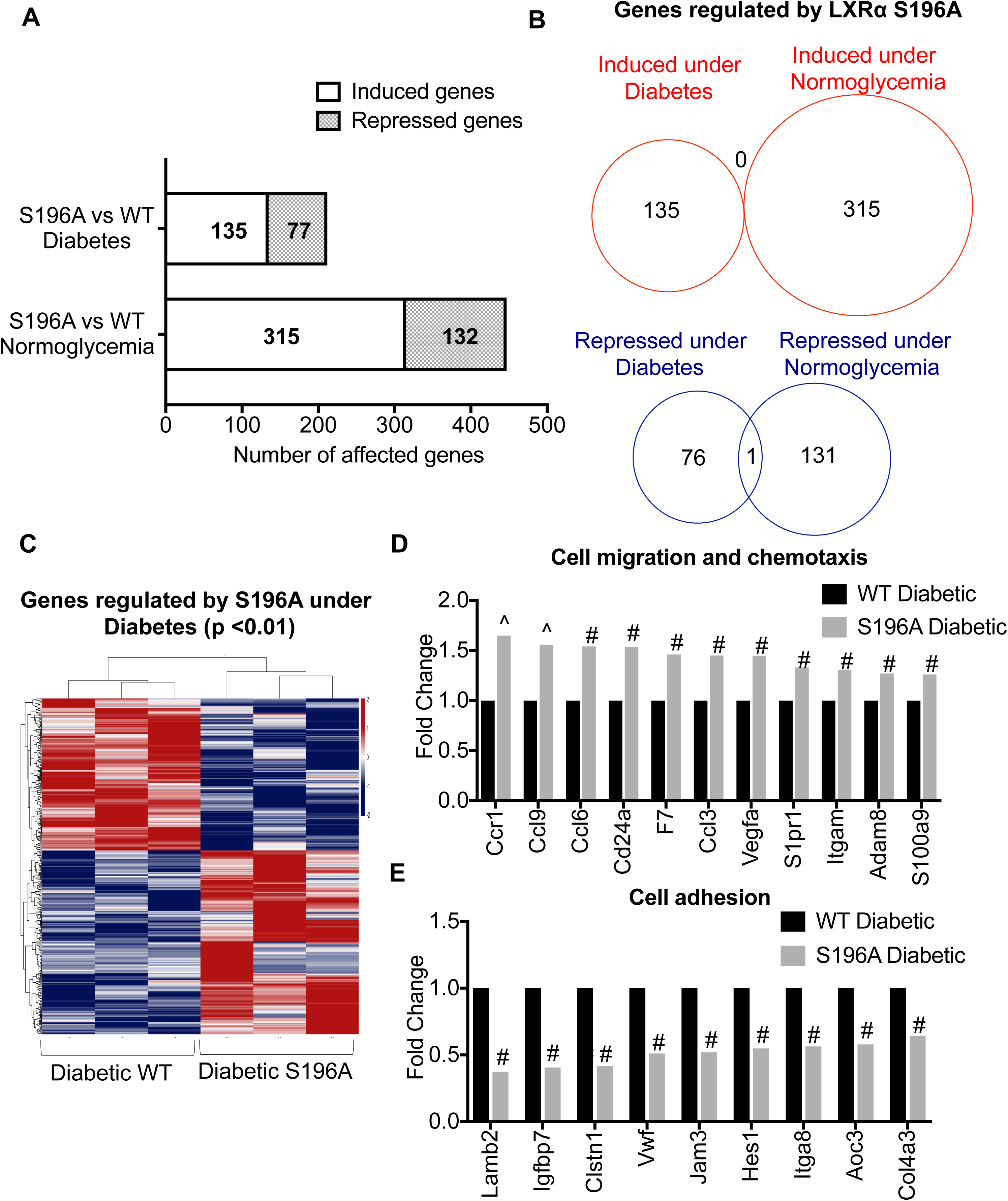
RNA-seq analysis reveals distinct set of genes regulated by LXRα S196A in plaque macrophages under normoglycemia and diabetes. A) Bar diagram showing the number of genes that exhibited >1.2-fold expression changes (p<0.01) (induced and repressed) from LXRα S196A under normoglycemia and diabetes. B) Venn diagrams showing overlap between genes regulated by LXRα S196A under normoglycemia and diabetes in plaque macrophages: induced genes are shown on the top panel and repressed genes on the bottom. C) Heatmap showing genes differentially expressed in LXRα S196A vs. WT groups under diabetic condition (P-value <0.01). D) Genes upregulated (>1.2 fold, # P-value <0.01, ^ P-value <0.001) by LXRα S196A under diabetic condition that are involved in cell migration and chemotaxis E) Genes downregulated (>1.2 fold, # P-value <0.01, ^ P-value <0.001) by LXRα S196A under diabetic condition that are involved in cell adhesion.

Genes upregulated by LXRα S196A under normoglycemia (i.e. repressed by LXRα pS196) (Supplementary Figure 10A) belonged to Gene Ontology (GO) classes that included cell adhesion, wound healing, cell differentiation and positive regulation of apoptotic process (Table 1). Genes downregulated by LXRα S196A (i.e. activated by LXRα pS196) aligned to GO classes that contained inflammatory response, chemotaxis, response to hypoxia and blood coagulation. This suggested that LXRα S196A expression promoted cell adhesion, and suppressed inflammation under normoglycemia. Moreover, a number of genes involved in positive regulation of apoptosis were upregulated by LXRα S196A consistent with increased apoptosis observed in the LXRα S196A plaques compared to LXRα WT under normoglycemia (Supplementary Figure 10B).

**Table 1.**
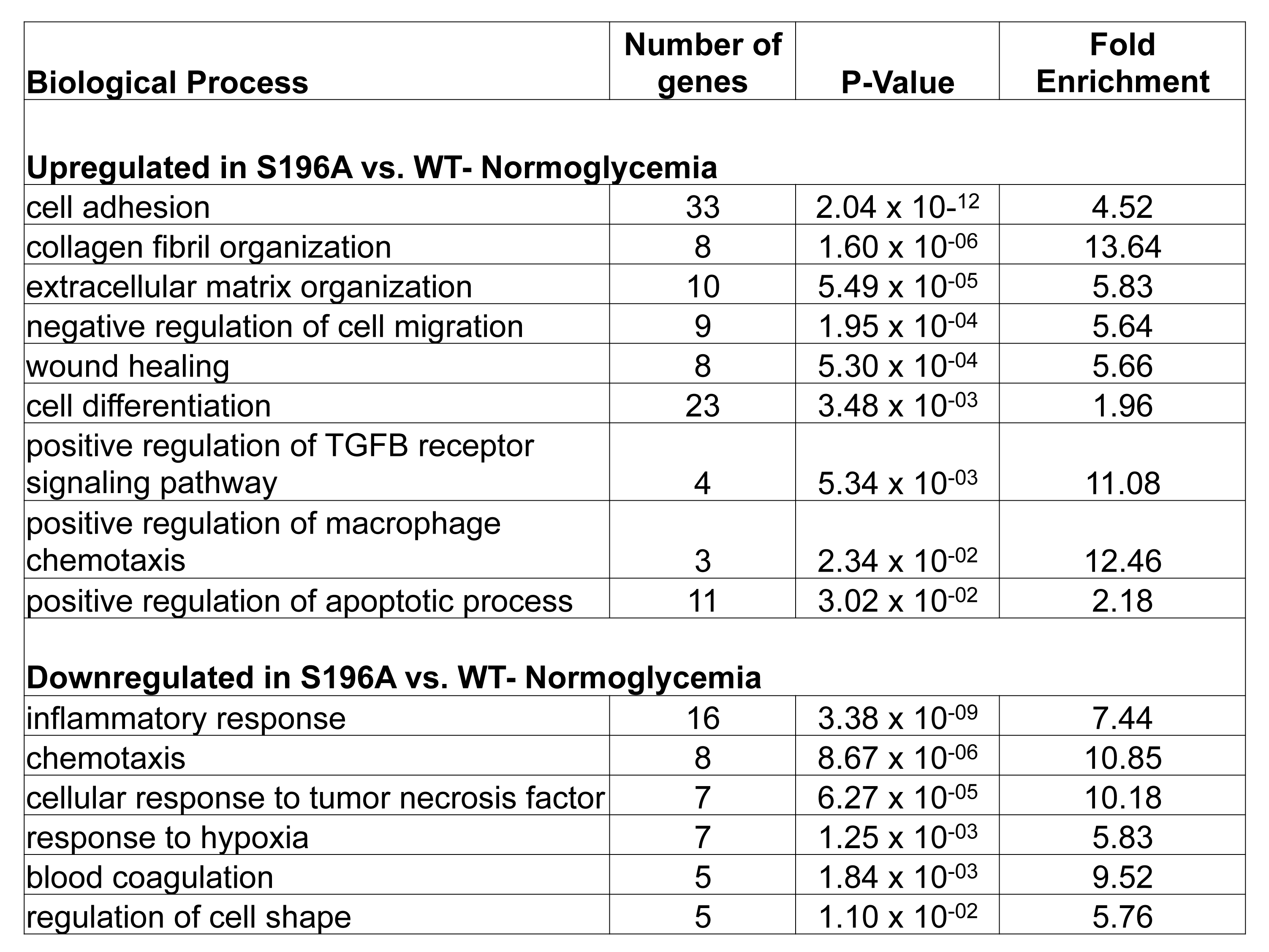
Gene Ontology analysis of genes upregulated or downregulated by LXRα S196A vs. LXRα WT expressing plaque macrophages under normoglycemia. Fold enrichment of actual over expected probesets along with the P-values are displayed for each biological process.

We next interrogated genes differentially expressed in LXRα S196A vs. WT under diabetic conditions (Figure 6C). GO classes enriched in genes upregulated by LXRα S196A (i.e. repressed by LXRα pS196) include leukocyte migration, chemotaxis, immune response and cholesterol homeostasis (Table 2). In contrast, the genes downregulated by LXRα S196A (i.e. activated by LXRα pS196) were associated with cell adhesion, negative regulation of cell migration, negative regulation of TGFβ signaling and neutrophil homeostasis (Table 2). A subset of genes upregulated by LXRα S196A have LXR binding sites in their regulatory regions suggesting direct regulation by LXRα (25), including CCR6, a chemokine receptor involved in monocyte migration in atherosclerosis (26), S100A8 and S100A9, inflammatory proteins secreted at increased levels from macrophages from diabetic mice (27). This suggests that LXRα S196A expression reduced plaque macrophage retention in diabetes by enhancing the expression of genes in macrophage migration and chemotaxis while repressing genes involved in cell adhesion making the macrophages less adhesive to the plaque (Figure 6D and E).

**Table 2.**
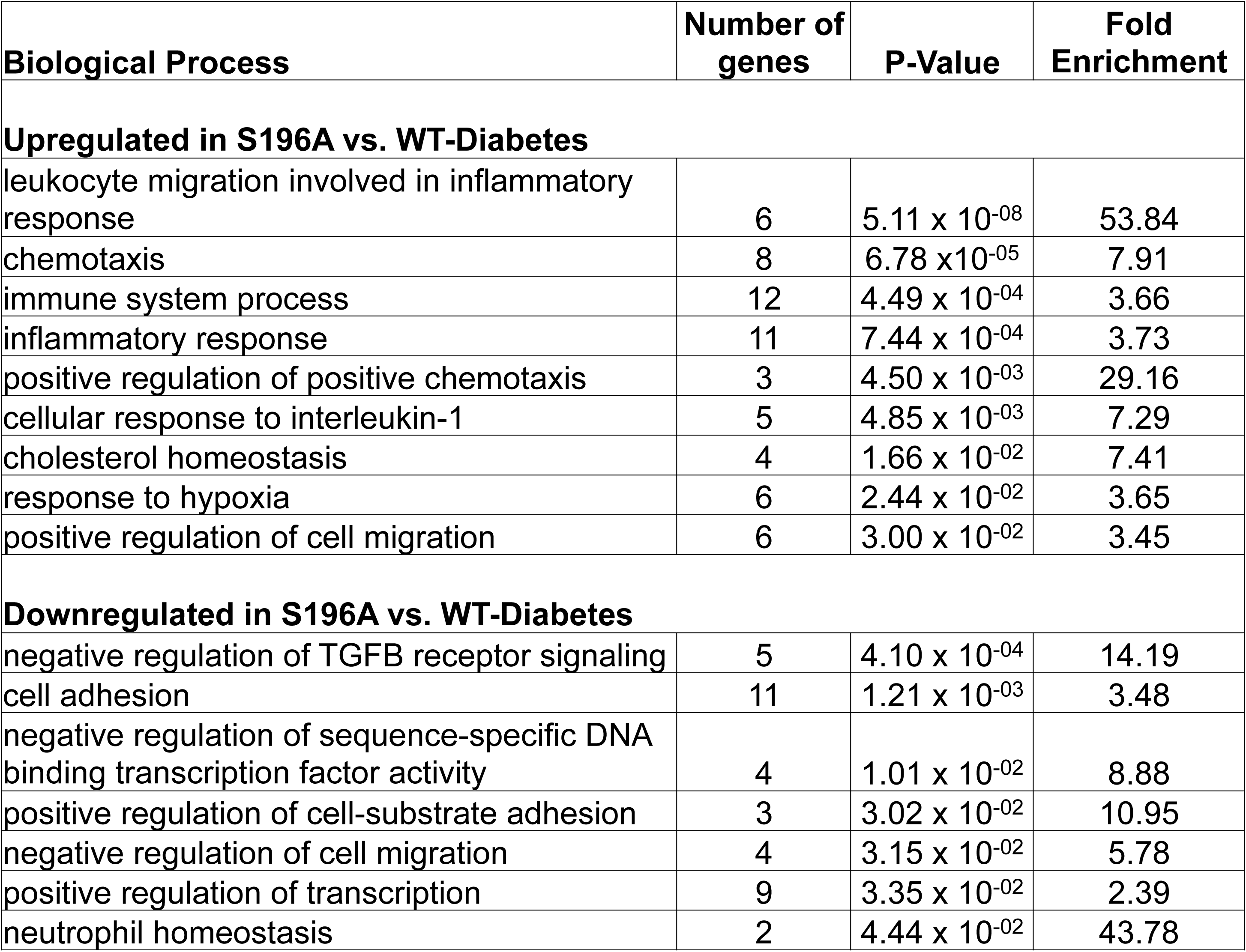
Gene Ontology analysis of genes upregulated or downregulated by LXRα S196A vs. LXRα WT expressing plaque macrophages under diabetes. Fold enrichment of actual over expected probesets along with the P-values are displayed for each biological process.

## Discussion

LXRα is post transcriptionally modified by phosphorylation (11, 26, 27), and yet the biological significance of this event is not well understood. In this study we examined the consequence of disrupting LXRα pS196 during regression of atherosclerosis in normal and diabetic mice. We found that diabetic mice expressing phosphorylation deficient LXRα S196A displayed reduced macrophage retention in plaques, suggesting the non-phosphorylated LXRα at S196 is anti-atherogenic under hyperglycemic conditions. We also found this occurs despite LXRα S196A promotion of leukocytosis through stimulation of bone marrow precursor proliferation (Figure 5C), which is expected to be pro-atherogenic (Figure 7). This promotion of leukocytosis was counterbalanced by the reduction of macrophage retention such that the plaques in S196A mice have similar macrophage content as in WT mice.

**Figure 7.**
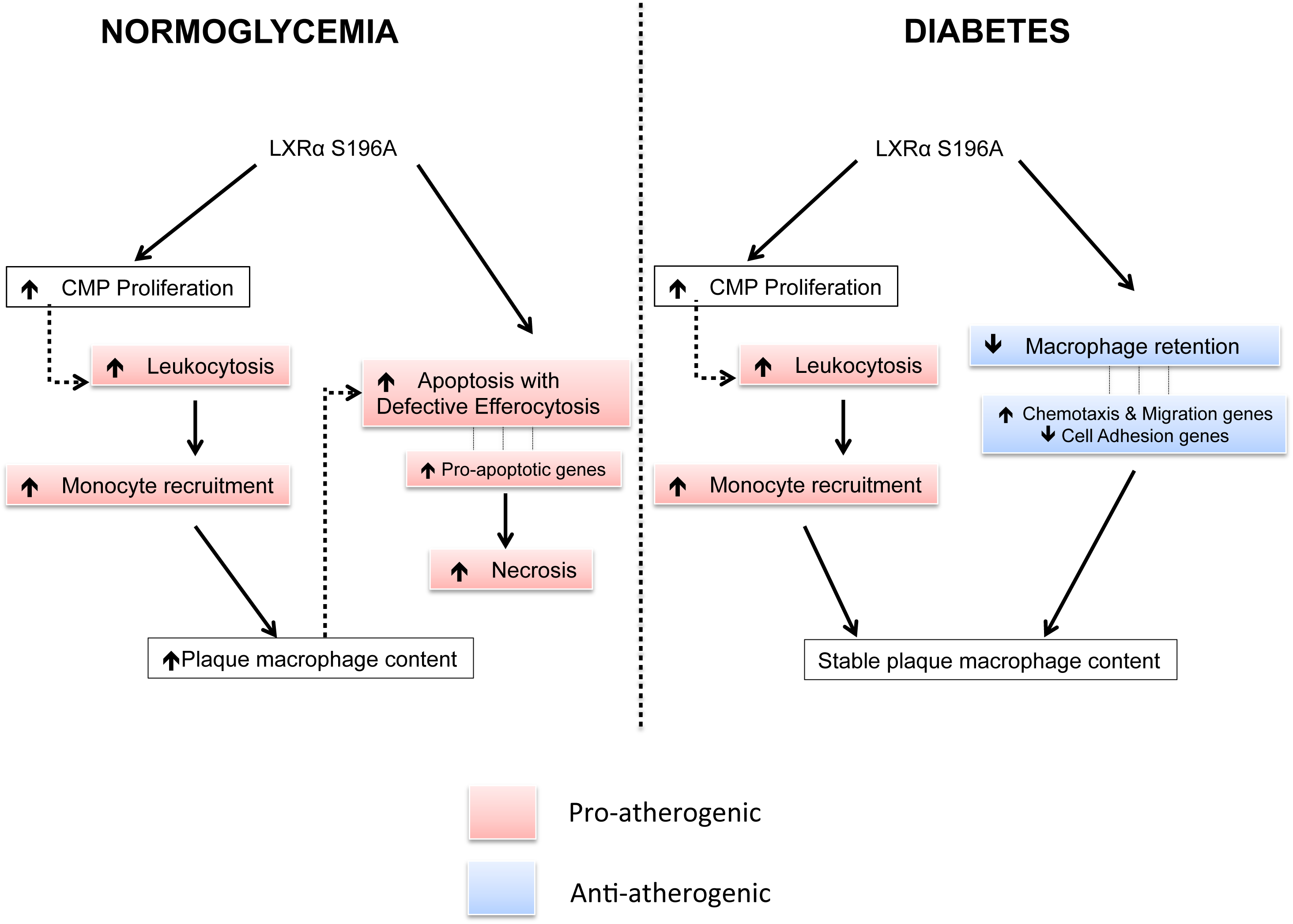
Effects of LXRα S196A expression under normoglycemia and diabetes in atherosclerosis regression. Under normoglycemic condition, LXRα S196A expression increases proliferating CMPs in the bone marrow that leads to increased circulating leukocytes. This increased monocyte infiltration, apoptosis and necrosis in the plaques leading to impaired regression under normoglycemia. Interestingly, LXRα S196A expression in diabetic mice leads to reduced macrophage retention and reverses macrophage accumulation in the plaque. This is through increased expression of genes promoting chemotaxis and migration, and decreased expression of genes involved with cell adhesion. This effect of LXRα S196A is anti-atherogenic. Therefore, despite the damaging effects of leukocytosis, LXRα S196A does not change plaque macrophage content and does not impair regression under diabetes compared to LXRα WT.

Phenotypes relevant to atherosclerosis such as plaque size, plaque necrosis and apoptosis, macrophage proliferation, plaque collagen content and macrophage efferocytosis were unchanged between LXRα WT and S196A groups under the diabetic setting. In contrast, plaque macrophage content, monocyte recruitment, apoptosis and necrosis were greater in LXRα S196A as compared to WT group under normoglycemia, likely as a result of increased monocytosis in S196A mice. Under diabetic conditions, there too was increased monocyte recruitment; however, this was mitigated by reduced macrophage retention in S196A compared to WT. This led to no change in plaque macrophage content between S196A and WT mice in diabetes. Thus, the phosphorylation state of LXRα modulates atherosclerosis regression dependent on different glycemic environments, via regulation of macrophage retention and the abundance of myeloid precursors.

Leukocytosis in LXRα S196A mice was unexpected, since there is no literature linking LXRα or its phosphorylation status to increased WBCs or monocytes. It has been previously shown that diabetes (19, 28), and reduced expression of the cholesterol transporters ABCA1 and ABCG1 in hematopoietic precursors, resulted in leukocytosis (29). We did not see changes in leukocyte numbers, however, as a function of diabetes. This might reflect mouse strain or methodological differences between studies. We also did not observe changes in ABCA1 and ABCG1 mRNA expression in plaque macrophages in S196A compared to WT mice; however, the regulation of these genes by LXRα S196A may differ in hematopoietic progenitor cells.

LXRα S198 phosphorylation was increased in RAW 264.7 macrophage cells expressing human LXRα when cultured in high compared to normal glucose. LXRα S198 has been reported to be phosphorylated by protein kinase A (PKA) (27) and CK2 (11), the later which exists as a tetramer with two α catalytic subunits (α and α’), and two regulatory β subunits. We did not see any upregulation in PKA or CK2 subunit mRNA expression by RNA-seq in macrophages from regressing plaques in normal versus diabetes. Nor did we see any change in CK2 α or β subunit protein abundance in the regressing plaques of normal versus diabetic mice by immunohistochemistry (not shown). This suggests that either there is post-translational activation of PKA or CK2 by hyperglycemia or the potential for other glucose sensing mechanisms to increase phosphorylation of LXRα, such as either decreased or increased activity of an LXRα pS196 phosphatase or kinase.

Differential gene expression analyses of macrophages (CD68+) in regressing plaques from WT versus S196A were consistent with the decreased macrophage retention or increased emigration in the plaques of S196A mice: the expression of genes involved in cell adhesion were decreased in the S196A compared to WT in diabetes while those in macrophage chemotaxis were increased. Strikingly, there was virtually no overlap in the gene expression signatures between WT and S196A plaque macrophages under normal or diabetes conditions. This could be due to alterations in LXRα occupancy genome-wide as a result of shifts in chromatin accessibility as a function of hyperglycemia. This could also be attributed to differences in the ability of the LXRα to interact with cooperating transcription factors under normal versus hyperglycemic conditions that promote receptor binding to target genes as a function of LXRα phosphorylation and thereby influence global gene expression (30). Thus, both LXRα phosphorylation and the cellular environment appear to play important roles in dictating gene expression in macrophages.

Taken together, our data suggest that phosphorylation deficient LXRα reduced the retention of macrophages in the plaque, and would therefore be atheroprotective under diabetic conditions. Harnessing the beneficial effects of suppressing LXRα phosphorylation systemically, however, would be problematic given the increased leukocytosis and enhanced monocyte recruitment to the plaque by the non-phosphorylated LXRα. To circumvent this issue, plaque-specific delivery of compounds that suppress LXRα phosphorylation or promote the conformation of the non-phosphorylated receptor might have therapeutic value in promoting the regression of atherosclerosis (31).

## MATERIALS AND METHODS

### Cell culture

RAW 264.7 cells stably expressing FLAG-tagged human LXRα previously described (11) were cultured in Dulbecco’s modified Eagle’s medium (DMEM, Corning) containing 10% fetal bovine serum (FBS) and 1 U/mL Penicillin and 1 μg/mL Streptomycin. Cells were routinely tested for mycoplasma and were mycoplasma free. Peritoneal macrophages were cultured in DMEM with 10% FBS. Jurkat cells were maintained in RPMI medium with 10% FBS with 1 U/mL Penicillin and 1 μg/ml Streptomycin.

### Microarray Analysis

Total RNA was isolated using RNEasy mini kit (Qiagen) and 3 μg of each sample was processed for hybridization to the mouse genome 430 2.0 AffyMetrix Expression Array by the Genome Technology Center of NYU Langone School of Medicine. Data are representative of four different conditions performed in quadruplicate and were normalized using Robust Multichip Average (RMA). The primary data were analyzed for fold-changes of >1.5 between treatment (T+9) and vehicle control cells (DMSO). Probesets were analyzed for significance (p < 0.05) using ANOVA. Genes were considered significant if they passed the ANOVA test and displayed a fold-change of >1.5 when the values between treated and control samples were compared. Ingenuity Pathway Analysis (IPA) software was used to analyze the gene sets for enriched molecular and cellular functions and canonical pathways.

### Preparation of whole cell extracts and immunoprecipitation

Cells were washed twice in PBS and lysed in Triton lysis buffer (50mM HEPES pH 7.6, 150mM NaCl, 1mM EDTA pH 8.0, 1mM EGTA pH 8.0, 1mM NaF, 1% Triton X-100, 10% glycerol) with protease and phosphatase inhibitor cocktail (Cell Signaling). Protein concentrations were measured by using the Bradford assay (Biorad). For immunoprecipitation, the lysates were incubated with FLAG antibody conjugated agarose beads overnight at 4°C. The beads were washed once in the lysis buffer and 3 times in TBS. Proteins associated with the antibody were eluted in TBS by competition with FLAG peptides.

### Western blotting

Protein samples were mixed with 5x Laemmli sample buffer containing beta-mercaptoethanol and boiled at 95°C for 5 min to denature the proteins. Proteins were separated on SDS-PAGE polyacrylamide gels and transferred to PVDF membranes (Millipore), incubated with blocking solution (5% BSA in TBS pH 7.4) for 1 h and incubated in primary antibody solubilized in blocking solution overnight at 4 °C. Antibodies used were anti-LXRα-pS196 (Affinity Purified 1135 (11)) and anti-LXRα (Abcam ab41902). Membranes were washed 3 times in TBS with 0.1% Tween (TBST) for 10 min and incubated with horseradish peroxidase-conjugated secondary antibody at room temperature for 1 h. Membranes were washed in TBST. The blots were developed with ECL detection reagents (E2400, Denville Scientific, Holliston, MA). Quantification of Western blots was done by using ImageJ software.

### Animals

Reversa mice (LDLR^−/−^; ApoB^100/100^; Mttp^fl/fl^; Mx1-Cre^+/+^) (19), global LXRα S196A KI mice, WT mice and inducible or non-inducible LXRα S196A phosphorylation deficient knock in mice (LXRα S196A^FL/FL^ RosaCreER^T2^ or LXRα S196A^FL/FL^ obtained from Dr. Ines Pineda, University College London, UK) in C57Bl6J background were cared for in accordance with the National Institutes of Health guidelines and the NYU Institutional Animal Care and Use Committee. Bone marrow from LXRα S196A^FL/FL^ or LXRα S196^FL/FL^ RosaCreER^T2^ mice were transplanted into 12 weeks old Reversa mice to generate mice with BM-derived, monocyte-derived cells capable of switching expression from LXRα WT to LXRα S196A upon tamoxifen treatment. After 8 weeks of BM transplantation, they were placed on Western diet (21% [wt/wt] fat, 0.3% cholesterol [Research Diets]) for 16 weeks to allow development of atherosclerotic plaques. At 14.5 weeks of western diet feeding mice were injected intraperitoneally (IP) daily with Tamoxifen (6 mg/40 g, Sigma-Aldrich) for 3 days to switch the expression from LXRα WT to LXRα S196A in LXRα S196^FL/FL^ RosaCreER^T2^ mice. After 2 days, mice were injected IP with STZ (50 mg/kg, Sigma-Aldrich) or citrate buffer for 5 days to induce diabetes mellitus or to serve as a control. Baseline groups were sacrificed at this time. The rest were then switched to a chow diet and were injected 4 times IP with polyinosinic polycytidylic RNA (Sigma-Aldrich 15 mg/kg) every other day (13). Regression groups were sacrificed after 5 weeks on chow diet. Mice were anesthetized with xylazine/ketamine, and blood was collected via cardiac puncture for plasma analyses. Mice were perfused with 10% sucrose/saline. Aortic roots were dissected and embedded in optimal cutting temperature compound medium (OCT) and frozen immediately and stored at −80°C until further use.

### Genotyping primers

The genotypes of S196A^FL/FL^, S196A^FL/+^, S196A^+/+^ mice were identified by PCR using 4 primers: R2: 5′−AAGCATGACCTGCACACAAG −3′; WT: 5′−GGT GTC CCC AAG GGT GTC CT −3′; F2: 5′−GGC ATG AGG GAG GAG TGT AA −3′; and SA: 5′−GGT GTC CCC AAG GGT GTC CG −3′. Primers R2 and WT identify the wild-type allele (642-bp) and FL allele (656-bp). Primers F2 and SA identify the FL allele (411-bp fragment). Primer R2 and SA identify mutant allele of LXRα S196A global knock in mice (656-bp). To genotype mice for RosaCreER^T2^ locus, Primers oIMR0092 [Mut], 5′−AAT CCA TCT TGT TCA ATG GCC GAT C−3′; oIMR3349 [common], 5′−CCA TAT GCA TCC CCA GTC TT−3′; and oIMR3350 [WT], 5′−GCG ATG GAT ACA CTC ACT GC−3′) were used. Primers oIMR0092 and oIMR3349 detect the mutant allele and oIMR3350 and IMR3349 detect the WT allele.

### Genomic DNA analysis in blood monocytes

Blood was collected from the mice after 8 weeks of bone marrow transplantation and after tamoxifen treatment and from S196A^FL/FL^, S196A^FL/+^, S196A^+/+^ mice as controls. Red blood cells were lysed with RBL buffer (Biolegend) and cells were stained with PECy7 anti-mouse CD45 and PE anti-mouse CD115 (Biolegend). Blood monocytes (CD45^hi^CD115^hi^) were sorted by FACS (Beckman Coulter MoFlo). Genomic DNA was extracted from the sorted monocytes using the DNEasy blood and tissue kit (Qiagen). Relative DNA levels of FL allele were quantified by qPCR using F2-SA primer pair. All the values were normalized to relative DNA levels detected using control primer pair. Primers used for DNA analysis are control DNA: forward 5’−AGG GAA AGC TCT CTG GAG CAT−3’ and reverse: 5’−CAG GAA TTT CTC CAT CCT TTG AGT−3’, F2-SA: forward 5’−GGC ATG AGG GAG GAG TGT AA−3’ and reverse 5’−GGT GTC CCC AAG GGT GTC CG−3’.

### Blood glucose and plasma lipoprotein analyses

Blood samples were centrifuged at 10,000 rpm for 10 minutes and the supernatants collected as plasma. Total cholesterol, triglyceride and HDL-C concentrations in plasma were measured using colorimetric assays (Wako Diagnostics, Richmond, VA). Blood glucose levels were measured with a blood glucometer (Freestyle lite, Roche) at the time of harvest.

### Histochemical analyses

Aortic roots frozen in OCT were serial-sectioned at a thickness of 6 μm onto glass slides. For CD68 (macrophage marker) staining, slides were fixed in 100% acetone for 10 mins and exposed to primary anti-CD68 antibody (Serotec), followed by biotinylated secondary antibody (Vector Laboratories), with visualization using a Vectastain ABC kit (Vector Laboratories). Sections were counter stained with hematoxylin. To stain for LXRα and pS196 in the plaques, sections were fixed in ice-cold acetone for 10 min, incubated in 3% H_2_O_2_ for 10 min to quench the peroxidase activity. Sections were then blocked with 5% normal goat serum (PBS) and stained overnight at 4°C with the primary antibody to LXRα (ab3585; Abcam) at 10 μg/ml and LXRα-S196-P (affinity-purified rabbit polyclonal 1135 (11)) in 5% normal goat serum-PBS. Sections were incubated with a secondary biotinylated anti-rabbit IgG (H+L) antibody (BA-1000; Vector Laboratories), followed by horseradish peroxidase streptavidin (SA-5704; Vector Laboratories). Staining was visualized using a DAB (3,3′-diaminobenzidine) peroxidase substrate kit (SK-4100; Vector Laboratories) following the manufacturer’s protocol, and the reaction mixture was counterstained with hematoxylin. Microscopic images were taken at 10X of aortic root sections and morphometric measurements were performed using Image Pro Plus software (Micro Optical Solutions).

### Apoptosis analysis

Apoptosis was analyzed in the aortic roots by staining the plaques with anti-cleaved caspase 3 (Cell signaling) and counter stained with hematoxylin. Nuclei positively stained for the antibody and hematoxylin were considered apoptotic. To quantify apoptosis in the plaque, % plaque cells (nuclei) positive for cleaved caspase 3 stain was calculated for each mouse.

### Proliferation analysis

Cell proliferation in the plaques were analyzed in aortic roots by staining the plaques for Ki67 (abcam). Nuclei positively stained for the antibody and hematoxylin were considered proliferating cells. To quantify proliferation in the plaque, the number of cells that stained positive for KI67 per section was measured for each mouse.

### Necrosis analysis

Necrosis was quantified in aortic roots by measuring the acellular areas of plaques with Image Pro Plus software as previously described (15).

### Laser capture microdissection

6 μm sections of aortic roots were collected on Pen membrane Frame Slides (Arcturus). CD68^+^ cells were isolated from atherosclerotic plaques by laser capture microdissection performed under RNase-free conditions (34, 35). Aortic root sections were stained with hematoxylin-eosin and cells were captured from approximately 36 frozen sections. After laser capture microdissection, RNA was isolated using the PicoPure Kit (Molecular Devices), and quality and quantity were determined using an Agilent 2100 Bioanalyzer (Agilent Technologies).

### RNA-Seq analysis

RNASeq libraries were prepared starting from 2.5 ng of total RNA from Baseline LXRα WT, LXRα WT Regression Control, LXRα S196A Regression Control, LXRα WT Regression STZ, LXRα S196A Regression STZ groups (n=3 or 4) using the Clontech SMARTer^®^ Stranded Total RNA-Seq Kit - Pico Input Mammalian, with 5 cycles of PCR for cDNA amplification and 13 cycles of PCR for library amplification, following the manufacturer’s protocol (Cat # 635006). Libraries were purified using AMPure beads, quantified by Qubit and QPCR, and visualized in an Agilent Bioanalyzer. The libraries were pooled equimolarly, and run on a HiSeq 2500, v4 chemistry, as paired end reads, 50 nucleotide in length. Raw sequencing data were received in FASTQ format and read mapping was performed using STAR 2.5 against the mm10 reference genome. The resulting BAM alignment files were processed using Featurecounts 1.0.5 on its respective GRCm38 annotation, obtained from the ENSEMBL database. Subsequently the Bioconductor package DESeq2 (3.4) was used to identify differentially expressed genes (DEG). This package provides statistics for determination of DEG using a model based on the negative binomial distribution. Genes with a p-value < 0.01 were determined to be differentially expressed.

### Monocyte/Macrophage tracking

For the egress study, Edu (1 mg/30 g) was injected IP at week 15 before STZ treatment to label Ly6C^hi^ monocytes, which can later become Ly6C^lo^ monocytes. For the recruitment studies, circulating Ly6C^lo^ monocytes were labeled by injecting mice in the retro-orbital vein with 250 μL 1μm Fluoresbrite fluorescein isothiocyanate-dyed (YG) plain microspheres (Polysciences Inc.) diluted in PBS (1:4) 24 hours before harvesting as previously described (36, 37). Labeling efficiency was assessed by flow cytometry 24 hours after injection. For EdU detection, blood samples were collected 24 hours and 5 days after injecting with EdU. Red blood cells were lysed with RBL buffer (BD Biosciences) and cells were stained with APC anti-mouse CD115, PeCy7 anti-mouse CD45 and PerCp-Cy5.5 anti-mouse Ly6G/C. EdU positive cells were detected with Click-iT^®^ EdU Flow Cytometry Assay Kit (Thermofisher, C10418), following manufacturer’s protocol. EdU positive cells in the plaques were detected using Click-iT EDU Imaging Kit (MP 10338) following manufacturer’s protocol. For bead detection, cells were stained with PECy7 anti-mouse CD45, PE anti-mouse CD115 and APC anti-mouse Ly6G/C. All antibodies were purchased from Biolegend. Bead labeling in cells was detected through FITC channel by flow cytometry and beads were counted in the plaques using a fluorescent microscope.

### Determining hematological parameters

Blood was collected before sacrificing the mice by tail bleeding. Total white blood cells (WBCs) and monocytes were determined by Genesis Hematology System (Oxford Science, Oxford, CT). Total monocytes were also calculated by multiplying total WBCs from Hematology system and % monocyte using flow cytometry analysis to confirm accuracy. Red blood cells were lysed with RBL buffer (company) and cells were stained with PECy7 anti-mouse CD45 and PE anti-mouse CD115 (Biolegend). Monocytes were identified by flow cytometry analysis as CD45^hi^CD115^hi^ cells using a LSRII analyzer.

### Hematopoietic stem and progenitor cells

Hematopoietic stem and progenitor cell populations were analyzed by flow cytometry. Bone marrow from S196^FL/FL^ or S196^FL/FL^ RosaCreER^T2^ mice were harvested 20 days after tamoxifen treatment, and red blood cells were lysed with RBL buffer (BD Biosciences). Cells were stained with a cocktail of biotinylated antibodies against lineage-committed (lin) cells (Gr1, B220, CD11b, CD3, TER119, CD8, CD4) and with antibodies for the stem and progenitor cell markers (Sca1, ckit, CD34, CD16/32, Cd150) Cells were identified as hematopoietic stem cells (HSCs; lin^−^, Sac1^+^, ckit^+^, CD34^-^, CD150^+^), common myeloid progenitor (CMP; lin^−^, Sca1^−^, ckit^+^, CD34^+^, CD16/32^mid^), and granulocyte–macrophage progenitor (GMP; lin^−^, Scal^−^, ckit^+^, CD34^+^ and CD16/32^hi^). Cell proliferation in each population was examined using DAPI by determining % cells in G_2_M phase (>2n DNA content). Flow cytometry was performed using an LSRII and all flow cytometry data were analyzed using FlowJo X software (Tree Star).

### Efferocytosis Assay

Peritoneal macrophages were isolated from mice by peritoneal lavage 3 days after injecting them intraperitoneally with 1 ml of 3% thioglycollate as previously described (38) and seeded in plated in a Lab-Tek Chamber Slide system (Thermo Scientific, 1256518) at a density of 2.5 x 10^5^ per well. Jurkat cells were labeled with Cell Tracker Red CMTPX dye (Molecular Probes, C34552) and rendered apoptotic by treating with cycloheximide (100 μg/mL) for 6 hrs. Apoptotic Jurkat cells in RPMI with 0.2 % BSA were added onto macrophages, at a 5:1 ratio of apoptotic cells: macrophages, and incubated for 2 hrs. Media was removed, and cells were washed with ice cold PBS three times to remove non-phagocytized apoptotic cells. Macrophages were fixed by incubating in 10% formalin for 10 minutes and washed with PBS three times. Macrophages were then stained with Phalloidin Green (Molecular Probes, A12379) and slides were mounted with DAPI containing mountant (Molecular probes, P36935). To quantify the extent of efferocytosis, macrophages with internalized apoptotic bodies were scored and results are expressed as % efferocytosis (number of macrophages with engulfed apoptotic cells / total number of macrophages x 100).

### Apoptosis Assay

Peritoneal macrophages were seeded in plated in Chamber Slide system as mentioned above. Apoptosis was induced by treating cells with LPS (100 μg/mL) and cyclohexamide (20 μg/ml) for 4 hrs. Cells were stained with FITC Annexin V and Propidium Iodide (BD Biosciences, 556547) for 15 minutes and washed 3 times with the binding buffer. Macrophages were fixed by incubating in 10% formalin for 10 minutes and washed with PBS three times. Slides were mounted with DAPI containing mountant (Molecular probes, P36935). Apoptotic cells (positive for Annexin V and negative for Propidium iodide) were scored and results are expressed as *%* apoptosis.

### Statistical analysis

The values are represented as means ± S.E. All comparisons were made from three independent experiments unless otherwise noted for all cell culture experiments. Statistical analyses were performed with Prism software (GraphPad Software, Inc., La Jolla, CA) using a 1-way ANOVA or a 2-way ANOVA or the two-tailed Student’s t test (^∗^, p ≤ 0.05; ^∗∗^, p ≤ 0.005; ^∗∗∗^, p ≤0.0005) as stated. Western blots are representative of three independent experiments.

## FIGURE LEGENDS

**Supplemental Figure 1. Signaling pathways affected by hyperglycemia in macrophages.** A) RAW 264.7 macrophages stably expressing human LXRα (RAW-LXRα) were cultured under high glucose (25mM D-glucose) or normal glucose (5.5mM D-glucose supplemented with L-glucose as an osmotic control) treated with vehicle (DMSO) or LXR ligands (5uM T0901317 +1uM 9cisRA) for 4 hours. Gene expression was assessed using the Mouse Genome 430 2.0 AffyMetrix Expression Array. B) Genes regulated by high versus normal glucose under ligand treated conditions were subjected to canonical signaling pathway analysis using IPA. Select signaling pathways predicted by IPA are shown.

**Supplemental Figure 2. Inducible phosphorylation deficient LXRα S196A knock in mouse genetics.** A) S196A^FL/FL^ mice are homozygous for Floxed (FL) allele which contains LXRα WT exon 5 (Ex5) and LXRα S196A exon 5 (Ex5’) of LXRα gene in opposite orientation flanked by lox sites sensitive to CRE recombinase. The mice express LXRα WT but can switch to LXRα S196A expression in the presence of CRE recombinase. We bred these mice to the Rosa26CreER^T2^ line that express CRE fused to Estrogen Receptor (ER) ligand binding domain in all cell types to obtain S196A^FL/FL^ RosaCreER^T2^ mice (homozygous for FL allele and heterozygote for Cre expressing allele). Administration of tamoxifen in these mice mediate recombination at lox sites resulting in the replacement of Ex5 with Ex5’. This allows Ex5’ to be in the proper orientation for expression, thereby switching expression from LXRα WT to LXRα S196A. B) DNA and amino acid sequences in LXRα WT (Ex5) and S196A (Ex5’) highlighting the serine to alanine mutation at amino acid 196.

**Supplemental Figure 3. Quantification of bone marrow reconstitution and tamoxifen induced CRE-mediated recombination efficiency** A) Schematic of a primer pair F2-SA that selectively amplifies FL allele in S196A^FL/FL^ mice. B) Blood monocytes were sorted by flow cytometry from S196A^FL/FL^ S196A^FL/+^ and S196A^+/+^ mice and total genomic DNA extracted. Relative FL DNA levels in S196A^FL/FL^ (arbitrarily set to 1), S196A^FL/+^ and S196A^+/+^ (WT homozygote) mice quantified by qPCR using F2-SA primer pair. C) Quantification of FL DNA levels in mice after BM transplant to monitor relative amount of BM donor derived FL DNA in Reversa mice receiving either S196A^FL/FL^ Cre- or S196A^FL/FL^ Cre+ BM (D) Quantification of FL DNA levels in mice after tamoxifen injections in S196A^FL/FL^Cre+ recipients to assess the CRE-mediated recombination efficiency.

**Supplemental Figure 4. LXRα pS196 abundance is reduced in plaques of S196A FL/FL Cre+ BM recipient compared to S196A ^FL/FL^ Cre-mice.** Aortic roots from the normoglycemic mice in the regression groups that were S196A ^FL/FL^ Cre+ or S196A ^FL/FL^ Cre-BM recipients were sectioned, fixed, and stained for LXRα pS196. Representative images of LXRα pS196 immunostaining are shown for each regression group.

**Supplemental Figure 5. Plaque total LXRα abundance is not altered by diabetes and S196A mutation.** Aortic roots from the mice in the regression groups were sectioned, fixed, and stained for LXRα. Representative images of LXRα immunostaining are shown for each regression group.

**Supplemental Figure 6. Triglyceride and HDLC levels.** A-B) Blood was collected from the mice at the time of harvest and plasma triglycerides (A) and HDLC (B) were measured by using colorimetric assays. Data (mean ± SEM) (n≥8) were analyzed using one-way ANOVA followed by Tukey’s multiple comparison test. P < 0.05 values were considered to be significant, ^∗^P < 0.05, ^∗∗^P < 0.005, ^∗∗∗^P < 0.0005 and ^∗∗∗∗^P < 0.00005. WT; S196A^FL/FL^ Cre-recipient Reversa mice; SA: S196A^FL/FL^ Cre+ recipient Reversa mice

**Supplemental Figure 7. Total plaque area is not affected by LXRα S196A expression.** The total area of the plaques was quantified using Image Pro Plus Software. Data (mean ± SEM) (n≥8) were analyzed using one-way ANOVA followed by Tukey’s multiple comparison test. P < 0.05 values were considered to be significant, ^∗^P < 0.05, ^∗∗^P < 0.005, ^∗∗∗^P < 0.0005 and ^∗∗∗∗^P < 0.00005. WT; S196A^FL/FL^ Cre-recipient Reversa mice; SA: S196A^FL/FL^ Cre+ recipient Reversa mice.

**Supplemental Figure 8. Fluorescent bead labeling and EdU incorporation in circulating cells were efficient and uniform across regression groups.** A) Reversa mice were injected with fluorescent latex beads 24 h before sacrifice to harvest aortic roots. Blood was collected prior to the harvest from the tail vein and precent circulating Ly6C^lo^ monocytes containing beads (FITC^+^) were measured by flow cytometry. B) Reversa mice were injected with EdU 6 days before the end of western diet feeding. To test the efficiency of EdU incorporation, blood was collected from LXRα WT and LXRα S196A expressing groups 24 h and 5 days after injecting EdU and percent EdU positive Ly6C^hi^ monocytes in circulation was measured by flow cytometry in both S196A^FL/FL^ RosaCreERT2 (WT) and S196A^FL/FL^ (S196A) recipient Reversa mice. C) The number of EdU positive cells was counted in aortic root plaques from the baseline groups. Data are expressed as mean ± SEM (n≥6 in each group). Data are expressed as mean ± SEM (n≥6 in each group). Data were analyzed using one-way ANOVA followed by Tukey’s multiple comparison test. P < 0.05 values were considered to be significant, ^∗^P < 0.05, ^∗∗^P < 0.005 and ^∗∗∗^P < 0.005.

**Supplemental Figure 9. LXRα S196A expression and diabetes do not alter plaque cell proliferation.** Aortic roots from the regression groups were sectioned, fixed, and stained for Ki67 to detect cells undergoing proliferation. The number of Ki67 cells per section was counted and are displayed in the graph. Data (mean ± SEM; n≥6) were analyzed using one-way ANOVA followed by Tukey’s multiple comparison test. P < 0.05 values were considered to be significant, ^∗^P < 0.05, ^∗∗^P < 0.005 and ^∗∗∗^P < 0.005.

**Supplemental Figure 10. Genes regulated by LXRα S196A under normoglycemic conditions** A) Heatmap showing genes differentially expressed in LXRα S196A vs. WT groups under normoglycemic condition (P-value <0.01). B) Genes upregulated (>1.2 fold, # P-value <0.01, ^ P-value <0.001) by LXRα S196A under normoglycemic condition that are involved in positive regulation of apoptosis.

**Supplementary Table 1.** Select genes regulated differently by LXRα S196A under normoglycemic and diabetic conditions (>1.2 fold, p<0.01). Fold-change compared to their LXRα WT counterpart under the given glucose setting are displayed with respective P-values.

## ACKNOWELDEMENTS

This work was support in part by NIH grant R01HL117226 (MJG, EAF), NIH training grant and Vilcek scholarship (ES), Medical Research Council New Investigator Grant G0801278 and British Heart Foundation Project Grant PG/13/10/30000 (IPT). We thank the NYU Genomics center for RNA-seq and bioinformatics support.

